# Does Balance Vary Across the Menstrual Cycle?: A Meta-Analysis with Implications for Women’s Health

**DOI:** 10.64898/2026.10.07.757426

**Authors:** Valeria Robles-Cerdas, Casey G. Turner, Christopher J. Brush, Julianne McGill, Abigail M. Skaarer, Niyati Baweja, Harsimran S. Baweja

## Abstract

**Background:** Women remain underrepresented across many domains of research, clinical science, and medicine. Sex differences other than the reproductive system, body composition, and size, are poorly understood. The menstrual cycle, for example, can be influenced using hormonal contraception methods. Hormonal fluctuations through the menstrual cycle go beyond the reproductive physiology and influence systems involved in balance and postural control.

**Data Sources and Methods:** The search was conducted through the following databases: PubMed, Web of Science, Medline, SPORTDiscus and Scopus (up to March 6, 2026) for studies that looked for the effects of the menstrual cycle phases on balance in women using or not hormonal contraceptives. Standardized mean difference was used to estimate the effect of menstrual cycle phases on postural control and Hedges’ *g* was reported to account for the small sample bias. Two multilevel mixed effects models, meta-regression, and sensitivity analyses were conducted. The funnel plot was used to determine if there was a publication bias. Thirteen studies were included in this meta-analysis.

**Results:** For static balance tasks, the pooled effect size was small and not statistically significant (SMDH = -0.19, 95% CI = -0.41 - 0.02, *p* = .073). Similarly, no significant effect observed for dynamic balance tasks (SMDH = 0.18, 95% CI = -0.67 - 1.04, *p* = .44). The moderator analyses showed that contraceptive use did not significantly explain the variability in either static or dynamic balance outcomes.

**Conclusions:** This meta-analysis demonstrated that current evidence does not establish a consistent effect of menstrual cycle phase on static or dynamic balance. High-quality studies with standardized menstrual cycle verification methods are needed to clarify the relationship between hormonal fluctuations and postural control in women.

**Registration:** PROSPERO CRD420261402114.

## Introduction

Women remain underrepresented across many domains of research, clinical science, and medicine (Abbasi, 2023; Becker et al., 2016; Sarkar & Sharma, 2026). This can lead researchers and the audience to miss important information about sex differences. Sex differences in balance and postural control, beyond those attributable to the reproductive system, body composition, and size, are poorly understood (Abo, 2022; Sims, 2018). However, the United States National Institutes of Health (NIH) established the Policy of Inclusion of Women and Minorities as Participants in Research Involving Human Subjects, requiring to include both women and men in all clinical trials and to report findings for both sexes (NIH, 2025). This has begun to narrow the gap in recent years, as different research fields have advanced our understanding of how the sex hormones can influence circadian rhythms, cancer therapeutics, cardiovascular diseases, and neuropathologies (Abo, 2022). Despite these advances, findings regarding the influence of hormonal fluctuations across the menstrual cycle on postural control remains unclear and inconsistently reported.

Sex differences in human physiology are driven by sex hormones and sex chromosome-dependent genetic mechanisms (Bale & Epperson, 2017). Women go through various stages throughout their lifespan, including the menstrual cycle, pregnancy, and menopause. Sex hormones influence multiple physiological systems, and their effects can differ across the menstrual cycle phases (Abo, 2022). Estrogen and progesterone are the primary sex hormones involved in the menstrual cycle. Previous research has shown that the hormonal fluctuations, including luteinizing hormone (LH), follicle-stimulating hormones (FSH), estrogen (estradiol), and progesterone, can be used to identify the different phases of the menstrual cycle (Gimunová, 2024). Levels of estrogen and progesterone are at their lowest during the menstruation phase, estrogen levels then rise during the follicular phase and continue increasing through the ovulation phase (Gimunová, 2024). After this, progesterone levels rise during the luteal phase. Finally, both hormone levels return to their baseline and initiate the next cycle (Abo, 2022). A regular menstrual cycle length is on average between 21 and 35 days (Gimunová, 2024).

The menstrual cycle is also influenced by the use of contraception methods. In the United States, during 2022 and 2023, 54.3% of females aged 15-49 reported using contraceptives. Among these, the two most common methods were the pill and long-acting reversible contraceptives, at 11.4% and 10.5% respectively (Daniels, 2025). It is important to emphasize that most of these methods are hormonal, suggesting that a considerable amount of women experience altered endogenous hormonal fluctuations (Ross & Eisenlohr-Moul, 2026).

Hormonal contraceptives can affect both reproductive and non-reproductive tissues. Previous research has identified estrogen and progesterone receptors in the uterus, ovaries, breast tissue, the pituitary gland, heart, muscles, and bones (El Khiati et al., 2023; Iorga et al., 2017; Simpson, 2003). Hormonal contraceptives act to suppress estrogen and progesterone levels by the inhibition of gonadotropin-releasing hormone secretion from the hypothalamus, which in turn prevents the pituitary gland from secreting follicle-stimulating hormone and luteinizing hormone (Sims, 2018). This suppression prevents the follicular development and ovulation, which leads to a decrease in the estrogen production and reduce progesterone levels due to the absence of corpus luteum, a temporary hormone-secreting gland produced in the ovaries after ovulation.

Hormonal fluctuation influences several systems involved in balance, including ligament laxity, muscle function and cognitive performance (Herzberg et al., 2017). Therefore, it is important to understand whether these hormonal changes affect postural control, as alterations in postural control may influence functional performance and increase the risk of falls and musculoskeletal injuries. Standing balance is the ability to maintain an upright position on two feet without falling (Goble, 2019). Postural sway, a natural movement, is a proxy measure of standing balance and helps to identify the individual’s risk of falling (Goble, 2018a, 2018b). Maintaining balance requires the interaction between the proprioception, vision, and vestibular inputs (Assländer L, 2014; Goble, 2025; Pasma JH, 2015). Because sex hormones can influence physiological systems that contributes to balance, hormonal fluctuations across the menstrual cycle may also alter postural.

Current evidence remains limited, and it is still unclear whether menstrual cycle phases significantly affect balance. Therefore, the purpose of this meta-analysis is to understand how phases of the menstrual cycle in eumenorrheic women (women with regular cycles) and women using hormonal contraceptives influence the ability to maintain balance. Improving our understanding of these relationships may help identify in which periods there is more susceptibility to impaired postural control. This could have implications for injury prevention, fall risk, and the development of individualized training and rehabilitation strategies for women across the lifespan.

## Methods

Static and dynamic tasks represent different aspects of balance; therefore, two separate meta-analyses were conducted. Both analyses used multilevel linear mixed effects models with Hedges’ g as the effect size metric. We considered for dependence among effect sizes within studies by using the study’s author as a random-effects and the total of effect sizes calculated nested within the study’s author.

### Search strategy

This meta-analysis was conducted in accordance with the Preferred Reporting Items for Systematic Reviews and Meta-Analysis (PRISMA) 2020 guidelines (Page et al., 2021). The search query was created by the lead author (VRC). The electronic search was executed by two reviewers (VRC and AS) through the following databases: PubMed, Web of Science, Medline, SPORTDiscus and Scopus. Search terms and Boolean operators were used in following format: ("Postural Balance"[Mesh] OR “postural balance”[tiab] OR “postural control”[tiab] OR “postural sway”[tiab]) AND ("Menstrual Cycle"[Mesh] OR “menstrual cycle”[tiab] OR menstruation[tiab] OR menses[tiab] OR "Contraceptive Agents, Hormonal"[Mesh] OR contracepti*[tiab]). Searched studies were eligible for inclusion if they were published in English and available up to March 2026. The search was conducted between February 18 until March 6, 2026. Search results were exported as Research Information Systems files (.ris) and imported into the Covidence Systematic Review tool. Duplicates were removed automatically and manually within Covidence. The electronic search, title and abstracts, and full text screening process were executed by two reviewers (VRC and AS). Full texts of the potential articles were screened by each reviewer. Articles that met the eligibility were included in this meta-analysis.

### Inclusion/Exclusion Criteria

Studies were included if they measured postural sway in eumenorrheic women, regardless of contraceptive use, and reported postural sway outcomes from at least two menstrual cycle phases. Studies were included whether balance task were performed in a unipedal or bipedal stance. Articles that reported neurological disorder, population under 18 or over 65 years old, and studies that did not report any menstrual cycle information were excluded from this meta-analysis. We included studies that have reported center of pressure (COP) displacement in static tasks such as standing in an upright position with eyes either open or closed. For dynamic tasks, studies reporting participant’s limits of stability were included. Sessions and balance tasks or assessments varied widely across studies (Table 1). Following the SAGER guidelines, eligibility was restricted to studies involving female participants because the purpose of this meta-analysis was to examine the relationship between menstrual cycle phases, hormonal contraceptive use, and postural control.

**Table 1.**
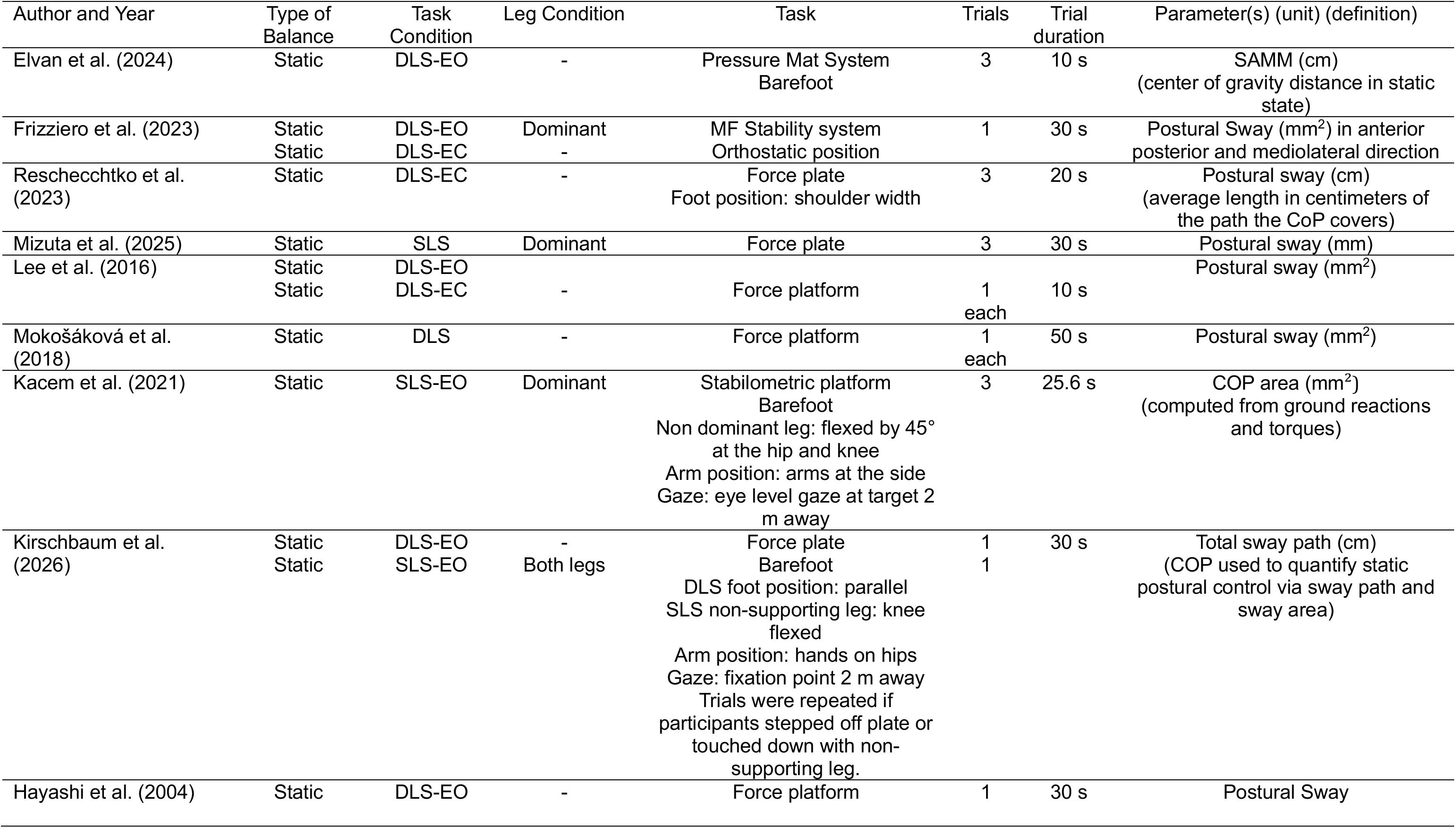
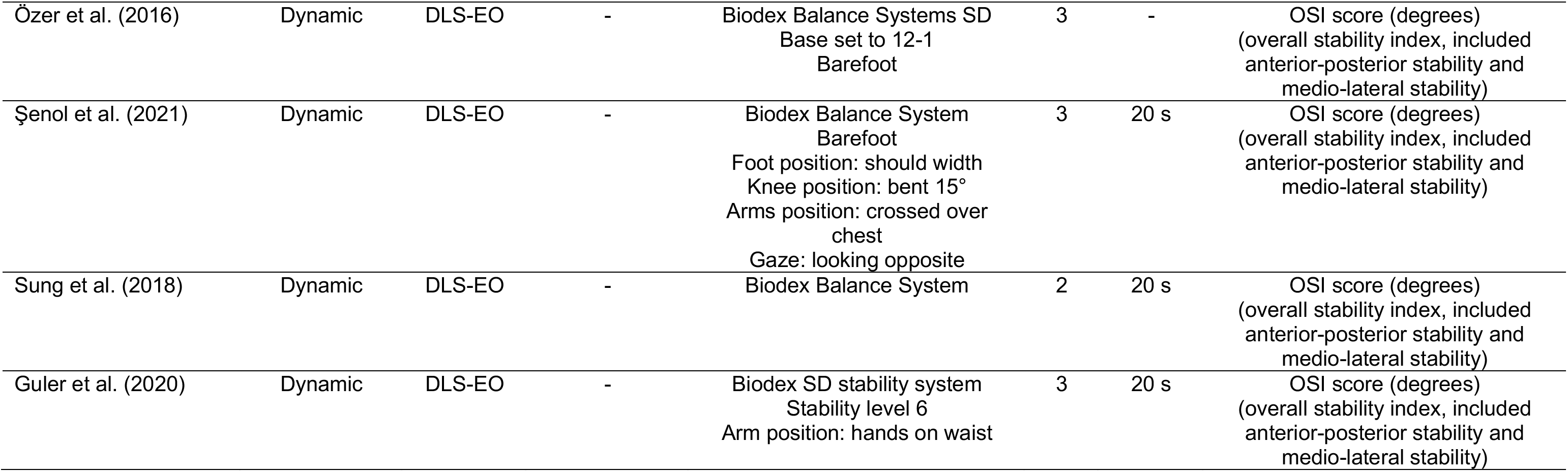
Methodological characteristics and experimental parameters of the included studies. Study descriptions detailed by author and year, type of balance (static or dynamic), task condition (DLS = double leg stance; SLS = single leg stance; EC = eyes closed; EO = eyes open), leg condition (dominant or non-dominant), task (balance assessment used), the amount trial performed and their duration, and the extracted balance outcome (postural sway or COP = center of pressure displacements)

| Author and Year | Type of Balance | Task Condition | Leg Condition | Task | Trials | Trial duration | Parameter(s) (unit) (definition) |
| --- | --- | --- | --- | --- | --- | --- | --- |
| Elvan et al. (2024) | Static | DLS-EO | - | Pressure Mat System<br>Barefoot | 3 | 10 s | SAMM (cm)<br>(center of gravity distance in static state) |
| Frizziero et al. (2023) | Static | DLS-EO | Dominant | MF Stability system | 1 | 30 s | Postural Sway (mm<sup>2</sup>) in anterior |
|  | Static | DLS-EC | - | Orthostatic position |  |  | posterior and mediolateral direction |
| Reschecchtko et al. (2023) | Static | DLS-EC | - | Force plate<br>Foot position: shoulder width | 3 | 20 s | Postural sway (cm)<br>(average length in centimeters of the path the CoP covers) |
| Mizuta et al. (2025) | Static | SLS | Dominant | Force plate | 3 | 30 s | Postural sway (mm) |
| Lee et al. (2016) | Static | DLS-EO |  |  |  |  | Postural sway (mm<sup>2</sup>) |
|  | Static | DLS-EC | - | Force platform | 1 each | 10 s |  |
| Mokošáková et al. (2018) | Static | DLS | - | Force platform | 1 each | 50 s | Postural sway (mm<sup>2</sup>) |
| Kacem et al. (2021) | Static | SLS-EO | Dominant | Stabilometric platform<br>Barefoot<br>Non dominant leg: flexed by 45° at the hip and knee<br>Arm position: arms at the side<br>Gaze: eye level gaze at target 2 m away | 3 | 25.6 s | COP area (mm<sup>2</sup>)<br>(computed from ground reactions and torques) |
| Kirschbaum et al. (2026) | Static | DLS-EO | - | Force plate | 1 | 30 s | Total sway path (cm) |
|  | Static | SLS-EO | Both legs | Barefoot<br>DLS foot position: parallel<br>SLS non-supporting leg: knee flexed<br>Arm position: hands on hips<br>Gaze: fixation point 2 m away<br>Trials were repeated if participants stepped off plate or touched down with non-supporting leg. | 1 |  | (COP used to quantify static postural control via sway path and sway area) |
| Hayashi et al. (2004) | Static | DLS-EO | - | Force platform | 1 | 30 s | Postural Sway |
| Özer et al. (2016) | Dynamic | DLS-EO | - | Biodex Balance Systems SD<br>Base set to 12-1<br>Barefoot | 3 | - | OSI score (degrees)<br>(overall stability index, included<br>anterior-posterior stability and<br>medio-lateral stability) |
| Şenol et al. (2021) | Dynamic | DLS-EO | - | Biodex Balance System<br>Barefoot<br>Foot position: should width<br>Knee position: bent 15°<br>Arms position: crossed over<br>chest<br>Gaze: looking opposite | 3 | 20 s | OSI score (degrees)<br>(overall stability index, included<br>anterior-posterior stability and<br>medio-lateral stability) |
| Sung et al. (2018) | Dynamic | DLS-EO | - | Biodex Balance System | 2 | 20 s | OSI score (degrees)<br>(overall stability index, included<br>anterior-posterior stability and<br>medio-lateral stability) |
| Guler et al. (2020) | Dynamic | DLS-EO | - | Biodex SD stability system<br>Stability level 6<br>Arm position: hands on waist | 3 | 20 s | OSI score (degrees)<br>(overall stability index, included<br>anterior-posterior stability and<br>medio-lateral stability) |

### Data extraction

Two reviewers extracted data (VRC and AS), creating columns with data sources within the Excel file, using information directly from the studies tables and methods section. We extracted demographic information (i.e.: age, weight, height, BMI, population), menstrual cycle phase (i.e.: menstruation, follicular, ovulation, and luteal) menstrual cycle length, physical activity, physical activity levels, and duration of the study information (i.e.: 1 or 2 months). For balance test/assessments we extracted postural sway length from the graphs, using WebPlotDigitizer (Rohatgi, 2024) tool, for the studies that did not report it in text. For studies with missing data, authors were contacted through email to request additional information. Authors were given 4 weeks to respond, studies for which the requested data were not provided within this timeframe were excluded from this analysis.

### Effect Size

To estimate the effect of menstrual cycles phases on balance across studies, standardized mean differences corrected for small sample bias (Hedges’g; SMDH) were calculated (Hedges & Olkin, 1985; Hedges, 1981). For static postural control tasks, the follicular phase was used as the reference phase because it was the most consistently reported condition across studies. Standardized mean differences were calculated relative to menstruation, ovulation, and luteal phases. For dynamic postural control tasks, the luteal phase was used as the reference phase because dynamic balance outcomes were consistently reported across this phase in the eligible studies. Consequently, standardized mean differences were calculated relative to menstruation, ovulation and follicular phases. Effect sizes were calculated using an independent-groups design based on reported summary statistics. The multilevel mixed-effect model accounted for dependence among multiple effect sizes contributed by the same study by including study author as random effect and effect sizes nested within study author. Even though pooled estimates were generated, confidence in the evidence remains limited because the small sample of studies included in the analysis and methodological variables were inconsistently reported across researchers. Analyses were conducted using the computer software JASP (Version 0.96.0; JASP Team, 2026).

### Moderator Analysis

To explore the potential sources of heterogeneity and the factors influencing the effect size, moderator analyses were conducted examining hormonal contraceptive use. For both static and dynamic task analyses, studies were categorized as “mixed” for the studies that included both user and non-users, “no use” for studies with only non-users and “use” for studies with only users. This analysis allowed us to assess whether contraceptive status explained variability in effect sizes observed between menstrual cycle phases. Although several study characteristics may influence balance outcomes, including menstrual cycle verification method, balance tasks characteristics, participant activity levels, and contraceptive formulation, these variables were not consistently reported across the included studies. Therefore, hormonal contraceptive-use status was the only study level characteristic reported with sufficient consistency to permit moderator analysis.

### Sensitivity Analysis

A sensitivity analysis was necessary to ensure the results are credible and robust (Nair, 2024). Several procedures were conducted, including asymmetry test, Egger’s test and Begg’s test and Cook’s distance to identify possible overly influential studies.

### Protocol registration

This meta-analysis protocol was registered on PROSPERO (International Prospective Register of Systematic Reviews) in June 2026 (CRD420261402114).

## Results

A total of 318 studies were found after the initial online search. After adding them all to Covidence Systematic Review tool, 123 duplicates were detected and removed. Thus, a total of 195 were screened for inclusion. After a title and abstract screening, 35 papers were moved to full-text review. Of these, thirteen papers remained eligible for the analysis. Figure 1 explains the PRISMA flow diagram of this selection process. Studies included in this meta-analysis were conducted between 2004 and 2026.

**Figure 1.**
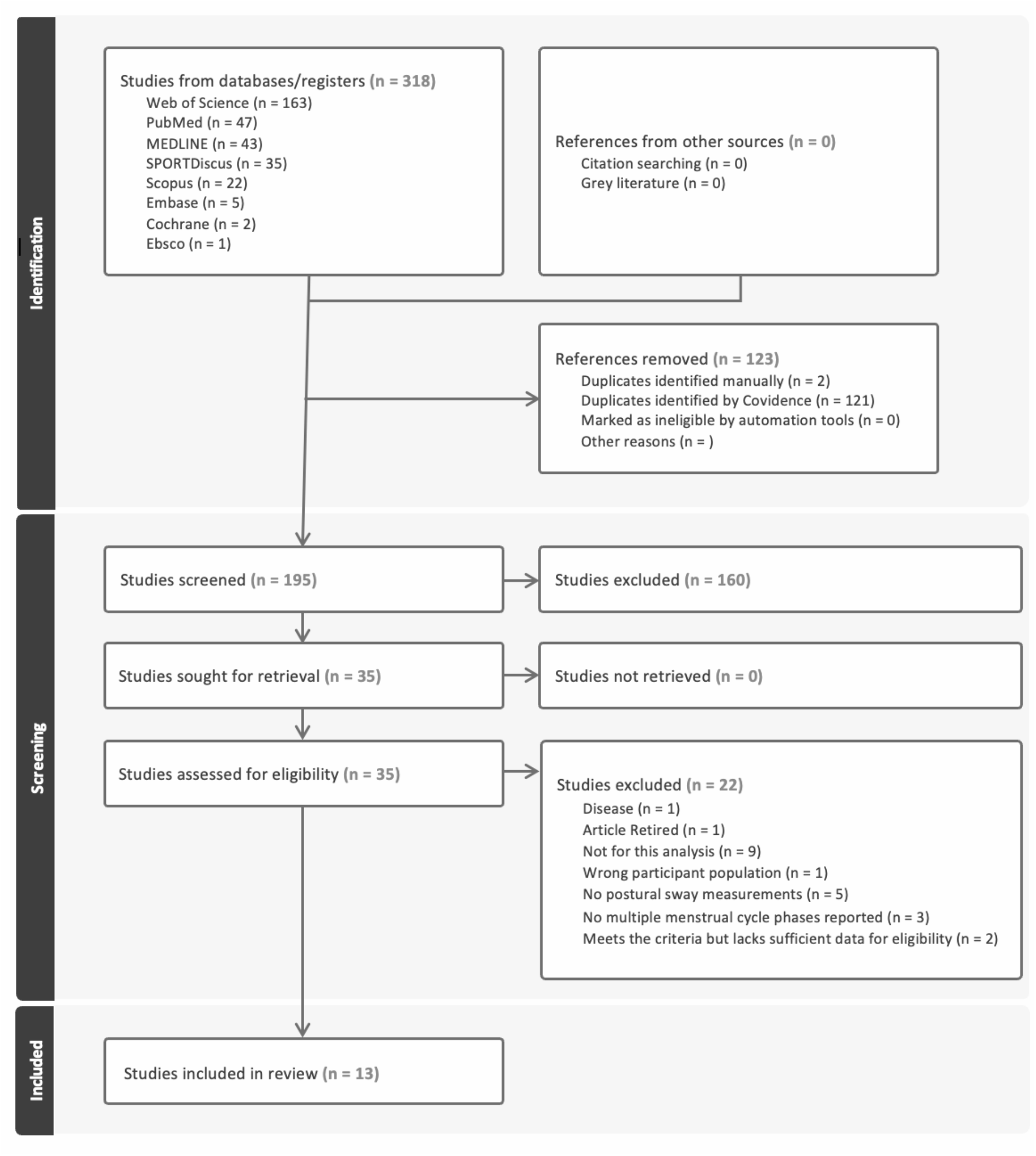
PRISMA 2020 flow diagram illustrating the study selection process for the systematic review and meta-analysis.

The methodological characteristics and experimental parameters of the included studies are summarized in Table 1. Studies varied in balance task type (static or dynamic), testing conditions, balance assessments, and outcome measures. Participant demographic and anthropometric characteristics are presented in Table 2, including sample size, contraceptive use status, and study-specific participant characteristics.

**Table 2.** Participants demographic and anthropometric characteristics across included studies. Studies are detailed by author and year, total sample size, status of contraceptive use, and specific study population criteria. BMI: body mass index; OC: oral contraception

| Author and year | Sample size | Contraceptive use | Population | Age (years) | Height (cm), weight (kg), BMI (kg/m<sup>2</sup>) | Cycle length (days) |
| --- | --- | --- | --- | --- | --- | --- |
| Elvan et al. (2024) | 63 | No use | Healthy women | 22±1.5 | -<br>58±7.4<br>21.3±2.36 | 30±3 |
| Frizziero et al. (2023) | 11 | No use | Healthy women | 29±3.41 | -<br>-<br>21.24±1.84 | 27-29 |
| Reschechtko et al. (2023) | 32 | Mixed | Healthy women | 21.34±1.73 | 163.78±7.65<br>61.02±7.65<br>- | - |
| Mizuta et al. (2025) | 33 | No use | Healthy women | 21.6±1.4 | 158.9±6.1<br>52±5.3<br>20.6±1.7 | 25-38 |
| Lee et al. (2016) | 14 | No use | Healthy women | 20.8±1.8 | 161.3±4.3<br>57.2±7.3<br>21.85±2.33 | - |
| Mokošáková et al. (2018) | 11 (control)<br>12 (OC) | Mixed | Healthy women | 24.64±1.54 (control)<br>23.33±0.92 (OC) | 171±2 (control)<br>168±2 (OC)<br>64±2.45 (control)<br>57.08±1.73 (OC)<br>21.75±0.88 (control)<br>20.23±0.33 (OC) | - |
| Kacem et al. (2021) | 12 | No use | Young healthy athletes | 21±1.6 | 172±0.05<br>65±5.6<br>- | 28±1 |
| Kirschbaum et al. (2026) | 6 | No use | Young healthy athletes | 25.8±3.3 | 171±6.8<br>68.1±6.4<br>25.8±3.3 | 21-35 |
| Hayashi et al. (2004) | 12 | No use | Healthy women | 20.4±1.2 | 159.1±4.7<br>50.4±2.8<br>20.4±1.2 | 25-35 |
| Özer et al. (2016) | 13 | No use | Healthy women | 21±1.25 | -<br>-<br>21.03±1.61 | 28.15±0.77 |
| Şenol et al. (2021) | 43 | No use | Healthy women | 20 | 166.5<br>20 | - |
|  |  |  |  |  | 20.8 |  |
| Sung et al. (2018) | 32 | No use | Healthy women | 19.83±0.92 | 160.67±6.53<br>58.5±6.67<br>22.69±2.54 | - |
| Guler et al. (2020) | 19 | Mixed | Healthy women | 20.47±1.07 | 164.78±4.66<br>54.63±3.41<br>20.11±0.97 | 28.47±1.26 |

### Meta-analysis: Static task

A total of 28 effect sizes from nine studies were included in the multilevel meta-analysis. The pooled effect size estimate comparing the follicular phase with non-follicular phases was -0.19 (95% CI: -0.41 - 0. 02), which was not statistically significant difference in balance outcomes across menstrual cycle phases, (*t*(_6.26_) = -2.15, *p* = .073). This pooled estimate represents the weighted average of all included studies. Influence diagnostics demonstrated that no individual study exerted a disproportionate effect on the model estimates, indicating that the overall findings were robust across studies.

### Moderator analysis: Hormonal Contraceptive Use

The moderator analysis demonstrated that contraceptive-use status was not associated with significant differences in effect size estimates, *Fm*_(2,1.54)_ = 8.64, *p* =.146. Specifically, studies involving naturally cycling women exhibited a pooled effect size estimate of -0.21 (95% CI = -0.47 - 0.05), whereas mixed samples demonstrated an estimate of 0.02, and hormonal contraceptives users demonstrated an estimate of -0.31 Although effect sizes varied numerically between groups, these differences were not statistically significant. Therefore, hormonal contraceptive use did not explain a significant proportion of the variability in effect size across studies.

Residual heterogeneity was not statistically significant after accounting for contraceptive-use status and the random-effects structure of the model (*Q*_e_(25) = 35.69, *p* = .077). The estimated between-study variance was 0.03, whereas the within-study variance (effect sizes nested within study) was negligible (σ^2^ = 6.875x10^-11^), indicating that most of the remaining variability occurred between studies rather than among effect sizes within studies.

### Meta-analysis: Dynamic task

A total of nine effect sizes from four studies were included in the multilevel meta-analysis. The pooled effect size estimate was 0.18 (95% CI: -0.67-1.04) which was not statistically significant (t(_1.88_) = 0.98, p= .44). These findings indicate that menstrual cycle phase was not associated with significant differences in dynamic postural sway across studies.

### Moderator analysis: Hormonal Contraceptive Use

The moderator analyses indicated that contraceptive use did not significantly influence the pooled effect estimate (*Fm*_(1, 1.92)_ = 17.21, *p* =.058). Studies involving naturally cycling women exhibited a pooled effect size estimate of -0.05 (95% CI = -1.14 - 1.05), whereas mixed samples demonstrated an estimated of 0.99 (95% CI = 0.04 - 1.94). Although effect sizes differed numerically between groups, the moderator effect did not reach statistical significance. Therefore, hormonal contraceptive use did not significantly explain variability in dynamic postural sway effect sizes across studies.

Residual heterogeneity remained significant after accounting for contraceptive-use status and the random-effects structure of the model (*Q*_e_(7) = 44.75, *p* < .001). The estimated between-study variance was negligible (σ^2 =^ 7.729x10^-10^), whereas the within-study variance (effect size nested within study) was substantially larger (σ^2^ = 0.63), indicating that most of the unexplained variability occurred among effect sizes within studies rather than between studies.

### Publication Bias and Sensitivity Analysis

To evaluate publication bias we conducted a sensitivity analysis. Figure 2A presents the funnel plot for static postural sway outcomes. Indicating an asymmetrical distribution, which was supported by significant Egger’s regression (*p* = .014) and Begg’s rank correlation (*p* = .013) test. The trim-and-fill analyses did not identify missing studies (*k*_0_ = 0), and the adjusted pooled effect estimate remain similar to the overall estimate. These findings suggest that the overall findings were relatively robust. Sensitivity analyses further demonstrated that no individual effect size exerted a disproportionate influence on the pooled estimate. The largest Cook’s distance value was 0.18, and standardized residuals remained small, indicating absence of extreme outliers.

**Figure 2.**
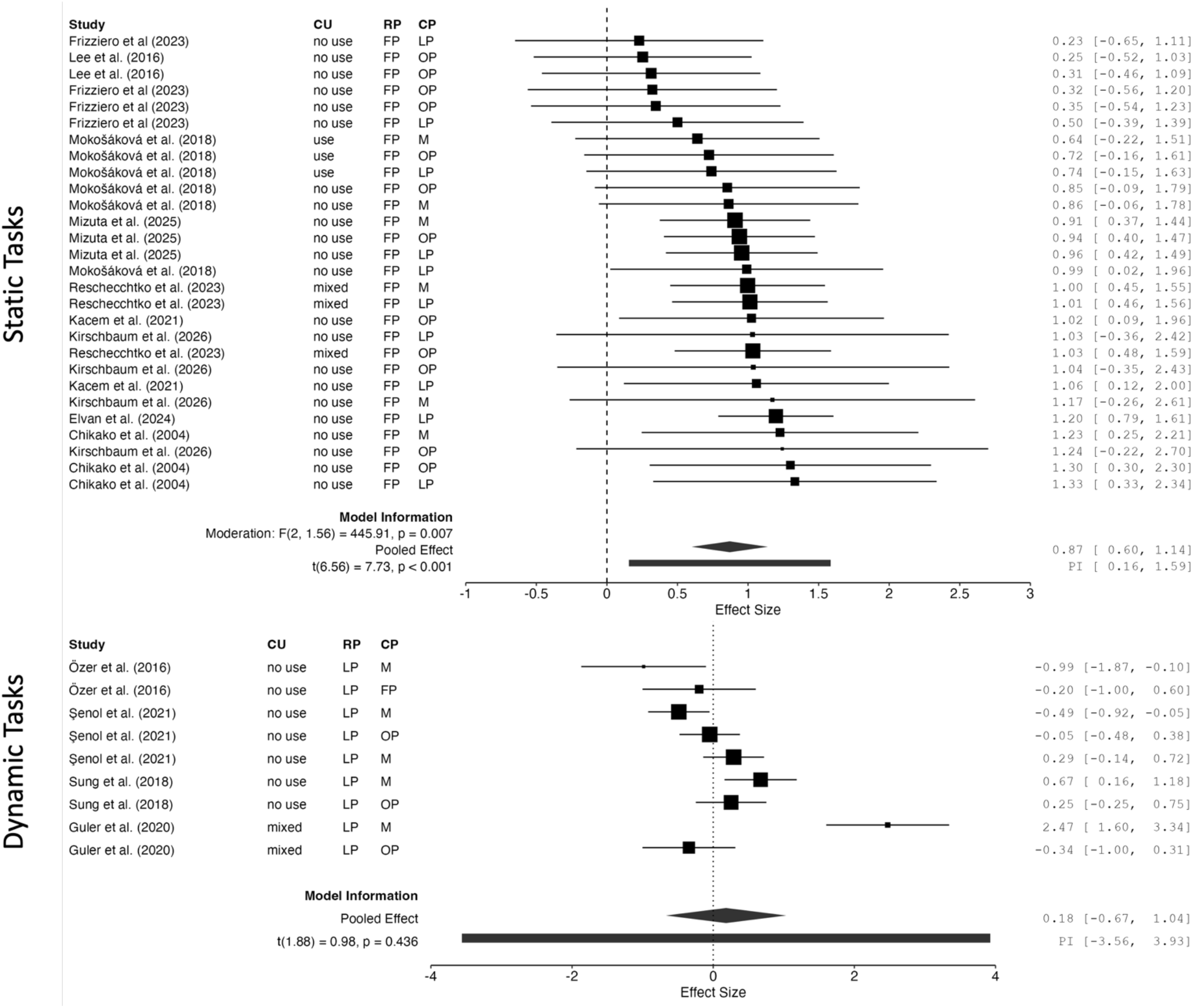
Static and Dynamic Tasks Forrest Plot. Each row represents one menstrual cycle phase (follicular) compared against other menstrual cycle phase (i.e.: menstruation, luteal, and ovulation). CU = contraceptive use, RF = reference phase; CP = comparison phase; FP = follicular phase; M = menstruation; LP = luteal phase, OP = ovulation phase. Each square represents the individual effect size estimates weighted; the diamond represents the pooled estimate and its 95% confidence interval.

**Figure 3.**
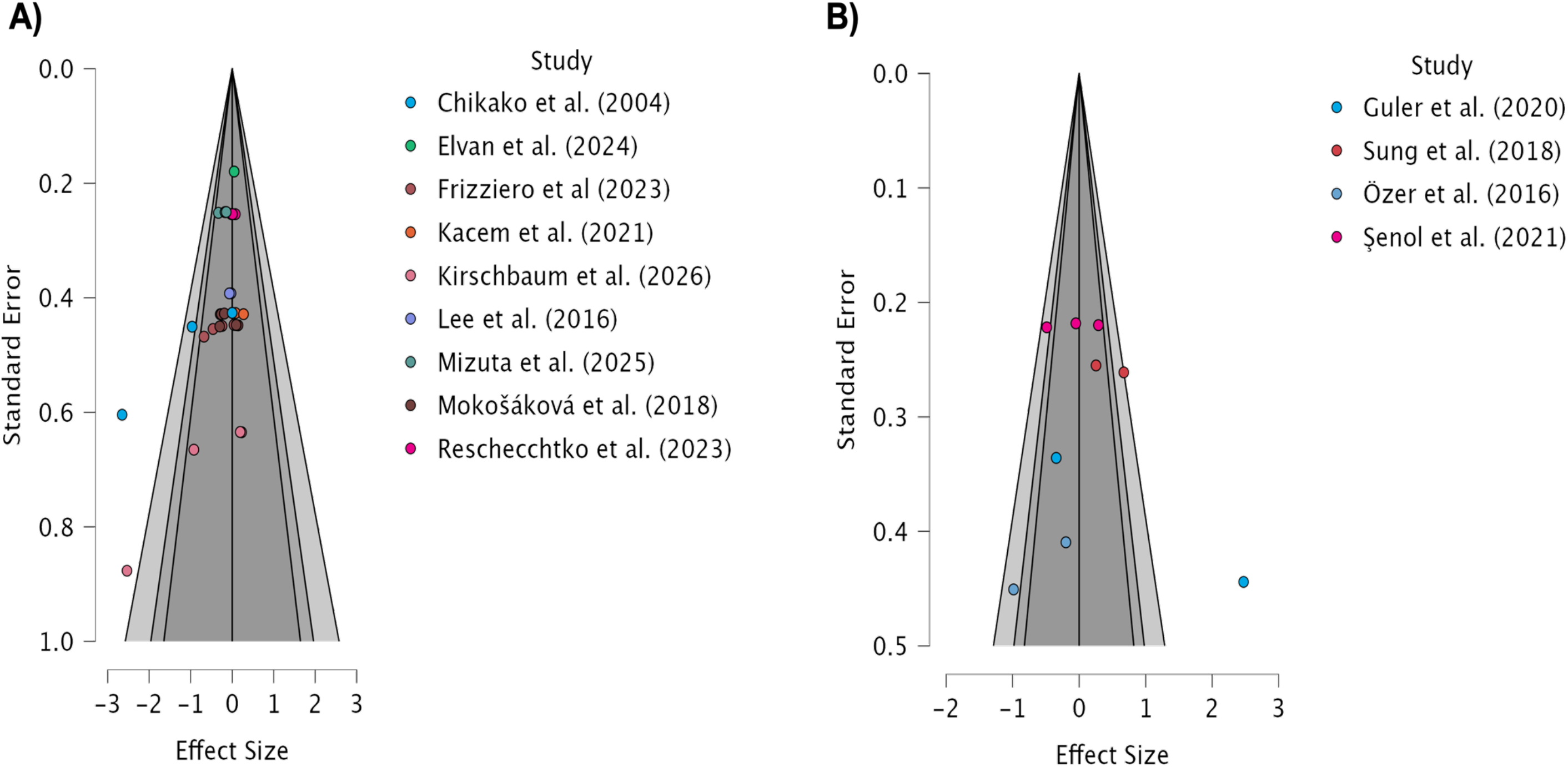
Funnel plots for publication bias assessment across static (3A) and dynamic (3B) tasks. Each data point represents on individual effect size estimate. The vertical axis plots the standard error as a function of the effect size on the horizontal axis. The dynamic tasks plot demonstrates a symmetric distribution of studies around the pooled estimate suggesting an absence of publication bias. However, the static tasks demonstrate an asymmetric distribution. The asymmetry results are reported in the text.

Figure 2B presents the funnel plot for dynamic postural sway outcomes. Although the distribution of effect sizes appeared relatively symmetrical, which was supported by a not significant Egger’s (*p* = 0.628) and Begg’s rank correlation (*p* = 0.919) test. Trim-and-fill analyses did not identify any missing studies (*k*_0_ = 0), and the adjusted pooled effect estimate remain non-significant. These findings provide no evidence of substantial publication bias and support the stability of the dynamic-tasks results. Given the small number of studies included, these findings should be interpreted with caution.

### Risk of Bias Assessment

The risk-of-bias assessment results for the thirteen included studies indicated that eight studies (61.5%) were judged as low overall risk of bias, four studies (30.8%) were judged as low overall risk and some concerns, and one study (7.7%) was classified as having some concerns. No studies were classified as high risk of bias. Across studies, low risk of bias was generally observed for participant selection, outcomes measurement, and statistical analysis domains. The most common concerns were related to menstrual cycle phase verification methods. Studies that relied solely on calendar-based phase estimation raised some concerns regarding the validity of menstrual cycle phase classification, whereas studies that incorporated hormonal assays, ovulation testing, basal body temperature monitoring, or multiple verification methods were rated as lower risk in this domain. Additional concerns were identified in some studies because of limited reporting of participant characteristics, contraceptive-use details, or potential confounding variables.

## Discussion

The purpose of this meta-analysis was to examine whether menstrual cycle phase and hormonal contraceptive use influence postural sway in women. Overall, our findings suggest that menstrual cycle phase was not associated with postural control either in static or dynamic balance tasks. Although the pooled effect estimates suggested a small numerical differences between phase, neither the static nor dynamic analyses reached statistical significance. These findings indicate that current evidence does not support a meaningful effect of menstrual cycle phase on postural sway across studies. We also found that hormonal contraceptive use produced numerically different effect sizes estimates across studies; however, these differences were not statistically significant. These findings suggest that differences in hormone exposure may help explain the inconsistent findings across studies.

The static tasks analyses demonstrated a small pooled effect size; however, this effect was not statistically significant. Fluctuations in estrogen and progesterone across the cycle have been shown to affect musculoskeletal tissues, ligament laxity, and central nervous system functions (Cabre, 2024; Chidi-Ogbolu, 2019). Nevertheless, the present findings do not provide sufficient evidence that menstrual cycle phase meaningfully alters static postural sway. Importantly, heterogeneity was not statistically significant, suggesting relatively consistent findings across studies, despite differences in study methods, participant characteristics, and outcome measures.

In contrast, the pooled effect for dynamic balance tasks was not statistically significant. Dynamic balance tasks involve movement and often rely on anticipatory and reactive control mechanisms. These mechanisms may make dynamic tasks less sensitive to subtle hormonal influences, or they may allow participants to compensate for changes in sensory or neuromuscular function (Massion, 1992). However, heterogeneity was statistically significant for the dynamic balance analysis, indicating meaningful variability across studies. This variability may reflect differences in study design, task demands, outcome measurements, and menstrual cycle phase definitions. Therefore, the current evidence in dynamic tasks may be not enough to determine whether menstrual cycle phases affect dynamic balance performance.

Hormonal contraceptive use is an important factor to consider when studying postural control in women. However, moderator analyses were not statistically significant for either the static or dynamic tasks. Although effect size estimates differed numerically among cycling women and hormonal contraceptive users, these findings do not support the conclusion that contraceptive-use status significantly explains variability in postural sway outcomes. Nevertheless, the numerical differences observed among groups suggest that hormonal exposure may warrant further investigation in future adequately powered studies.

Hormonal contraceptives suppress endogenous ovarian hormones fluctuations and alter the typical cyclical pattern of estrogen and progesterone exposure. These alterations may influence the physiological mechanisms involved in postural control. However, most studies included in this meta-analysis did not provide detailed information about contraceptive characteristics. This is an important limitation because hormonal contraceptives vary considerably in hormonal composition, dosage, administration route, and duration of use. Because these details were inconsistently reported, we were unable to examine specific contraceptive types or formulation separately.

Several strengths of this study should be acknowledged. First, the included studies represented women from several countries, including the United States, Turkey, Italy, Japan, Republic of Korea, Slovakia, Tunisa, and Germany. This international representation improves the breadth of the evidence base. Second, the use of multilevel meta-analytic methods allowed us to include multiple effect sizes from the same study while accounting for statistical dependence among outcomes. This approach allowed for a more comprehensive synthesis of the available evidence

Despite these strengths, several limitations should be considered. The number of eligible studies was small, particularly for the dynamic balance outcomes and contraceptive-use comparisons, which may have limited statistical power. Across studies, there was also variability in menstrual cycle phase definitions, phase verification methods, balance tasks, and postural sway outcomes. In addition, most studies did not report key contraceptive details, including type, formulation, dosage, route of administration, or duration of use. This limited our ability to examine whether specific contraceptive characteristics influenced postural control. Most studies assessed the same participants across multiple menstrual cycle phases. However, the information required to calculate repeated-measure effect sizes, was not consistently reported. This may have reduced the precision of the variance estimates.

Finally, this meta-analysis focused on postural sway outcomes and did not directly evaluate sensory reweighting, vestibular function, proprioception, or other mechanisms that may explain changes in balance. The ability to investigate sources of heterogeneity was limited by incomplete reporting within the included studies. Potentially important variables, including menstrual cycle phase verification method (e.g., calendar tracking, luteinizing hormone tests, hormones level confirmation), contraceptive formulation and duration of use, participant physical activity level, and balance testing procedures were inconsistently reported. As a result, these variables could not be use as moderators. Future studies should standardized reporting practices to improve the comparability of findings and facilitate more comprehensive research.

Additionally, studies are also needed to examine dynamic balance, sensory integration, and postural control across different stages of the female lifespan, including adolescence, perimenopause, and post menopause. These efforts will help clarify how hormonal fluctuations influence balance and whether these effects have meaningful implications for injury risk, rehabilitation, and physical performance.

## Conclusions

This meta-analysis examined whether menstrual cycle phases and hormonal contraceptive use influence postural sway in static and dynamic tasks in women. Despite a growing interest in women’s health and the potential influence of hormonal fluctuations on physical function, the current evidence does not indicate that menstrual cycle phase has a consistent or clinically meaningful impact on postural sway. Despite that the individual studies have reported phase-related differences in balance; these effects were not consistently supported when findings were synthesized across studies. Suggesting that any influence of menstrual cycle phase on postural control may be small, context-dependent, or obscured by methodological differences across studies. Similarly, while hormonal contraceptive use remains an important consideration in women’s health research, the available evidence was insufficient to demonstrate that contraceptive-use status meaningfully explains differences in postural sway across the menstrual cycle. The numerical differences observed between groups may indicate potential hormonal influences, but current evidence does not allow firm conclusions regarding their magnitude or direction.

Taken together, these findings highlight the need for more rigorous and standardized research in this area, particularly with respect to menstrual cycle phase verification, contraceptive-use reporting, and balance assessment protocols. Future studies should also investigate the underlying mechanisms that may link hormonal fluctuations to postural control, including sensory integration, vestibular function, proprioception, and cognition.

From a clinical perspective, the present evidence does not support modifying balance assessment or rehabilitation practices based solely on menstrual cycle phase or hormonal contraceptive use. Nevertheless, documenting these factors may improve the interpretation of individual responses and contribute to a more comprehensive understanding of women’s health across the lifespan. This may be especially relevant for individuals whose function or injury risk depends heavily on postural stability, including athletes, older women, and individuals recovering from lower-extremity injuries.

## Declaration of Conflicting Interests

### Authors’ Contributions

CrediT: **Valeria Robles-Cerdas**: Conceptualization, methodology, literature search, project administration, data extraction, formal analysis, writing - original draft, writing - review & editing. **Casey G. Turner**: Conceptualization, supervision, methodology, writing - review & editing. **CJ Brush**: Conceptualization, supervision, methodology, writing - review & editing. **Julianne McGill**: Conceptualization, supervision, methodology, writing - review & editing: **Abigail Skaarer:** Literature search, data extraction, methodology, writing - review & editing. **Niyati Baweja:** Methodology, writing - review & editing; **Harsimran S. Baweja**: Conceptualization, methodology, literature search, project administration, data extraction, formal analysis, writing - original draft, writing - review & editing.

### Data Availability Statement

Data supporting the findings of this study are available from the corresponding author upon reasonable request.

### Ethical Approval

Ethical approval was not required for this study because it involved analysis of previously published literature and did not involve human participants or identifiable personal data.

### Use of Generative Artificial Intelligence (AI)

A Generative AI tool (Claude Opus 5, Anthropic, PBC, San Francisco, CA, USA) was used solely in an assistive capacity, to support language editing, grammar, and readability. No generative AI tool was used to generate scientific content, to conduct or interpret analyses, or to produce references, tables, or figures. All text was reviewed and verified by the authors, who take full responsibility for the accuracy, originality, and integrity of the manuscript in its entirety.

### Abbreviations

NIH: National Institutes of Health
COP: Center of pressure
BMI: Body mass index
CI: confidence interval
DLS: double leg stance
SLS: single leg stance
EC: eyes closed
EO: eyes open
CU: contraceptive use
RF: reference phase
CP: comparison phase
FP: follicular phase
M: menstruation
LP: luteal phase
OP: ovulation phase

